# Detection of HPV E7 transcription at single-cell resolution in epidermis

**DOI:** 10.1101/252858

**Authors:** SW Lukowski, ZK Tuong, K Noske, A Senabouth, QH Nguyen, HP Soyer, IH Frazer, JE Powell

**Author notes:** These authors contributed equally (SW Lukowski, ZK Tuong, IH Frazer, JE Powell).

## Abstract

Persistent human papillomavirus (HPV) infection is responsible for at least 5% of human malignancies. Most HPV-associated cancers are initiated by the HPV16 genotype, as confirmed by detection of integrated HPV DNA in cells of oral and anogenital epithelial cancers. However, single-cell RNA-sequencing (scRNA-seq) may enable prediction of HPV involvement in carcinogenesis at other sites. We conducted scRNA-seq on keratinocytes from a mouse transgenic for the *E7* gene of HPV16, and showed sensitive and specific detection of HPV16-*E7* mRNA, predominantly in basal keratinocytes. We showed that increased *E7* mRNA copy number per cell was associated with increased expression of E7 induced genes. This technique enhances detection of viral transcripts in solid tissue and may clarify possible linkage of HPV infection to development of squamous cell carcinoma.

## Introduction

Squamous cell carcinoma (SCC), a non-melanoma keratinocyte malignancy, is one of the most common, potentially life-threatening cancers, and presents a major health care burden (Boukamp, 2005; Franceschi et al., 1996; Gordon et al., 2017). The risk of developing SCC is typically associated with genetic pre-disposition, increased exposure to sun and UV, and immunosuppression (Wheller and Soyer, 2015). SCC can also arise from actinic keratosis (AK) and intraepidermal carcinoma (IEC), both non-invasive precursor conditions (Feldman and Fleischer, 2011; Zalaudek et al., 2012). In immunocompetent patients, AK lesions commonly regress (Criscione et al., 2009), with up to 70% of lesions regressing without intervention. Regression is enhanced if sun block is used (Gordon et al., 2009), which is consistent with the amelioration of UV induced local immune suppression (Yu et al., 2014); although, persistent lesions often progress to SCC over time (Foote et al., 2001). AK lesions and SCCs are more common in immunocompromised subjects (Madeleine et al., 2017), and in these subjects, AK regression likelihood is reduced while progression to SCC is increased. Papillomaviruses (HPVs) are epitheliotropic viruses, and persistent HPV infection is responsible for at least 5% of human epithelial cancer (Torre et al., 2015). While the vast majority of these cancers are squamous cell carcinomas (SCCs) that arise within the anogenital tract and oropharynx, and are associated with infection by “high-risk” mucosotropic alpha HPVs (α-HPVs) such as HPV16, cutaneous beta HPVs (β-HPVs) also can promote SCC development in the skin. The clearest evidence for the role of cutaneous β-HPVs in SCC development is the well-accepted association of “high-risk” β-HPVs, such as HPV8, with SCCs in patients with *epidermodysplasia verruciformis* (EV) (Gewirtzman et al., 2008). EV patients carrying a homozygous mutation in either *EVER1* or *EVER2* develop disseminated benign skin lesions that can progress to SCC over time and these are thought to be initiated by β-HPVs. Detection of β-HPV viral DNA has been associated with some cutaneous SCCs, although swabs from the top of tumors have been shown to have higher levels of HPV DNA compared to samples taken from within the lesions (Forslund et al., 2004). Furthermore, the presence of HPV DNA may not correlate with active transcription of its RNA (Arron et al., 2011). This has led to the hypothesis that the progression towards skin cancer does not require the HPV virus for the maintenance of malignancy (*i.e.* hit and run carcinogenesis (Pfister, 2003)). This is in contrast to cancers caused by high-risk α-HPVs, in which the viral genome is maintained, and viral genes contribute to the maintenance of the cancer phenotype. Nevertheless, there are strong correlations between the presence of β-HPVs in the skin and the incidence of SCC in organ transplant recipients undergoing immunosuppressive drug therapy (Genders et al., 2015), and between the burden of β-papillomavirus infection and the risk of subsequent development of SCC (Farzan et al., 2013). These data are supportive of a role of β-HPVs in cutaneous SCCs in a broader context, however the hit and run carcinogenesis hypothesis for β-HPVs in cutaneous SCCs has been challenging to prove. The association of β-HPV infection with SCC and AK therefore needs further research, as vaccination to prevent infection with β-HPVs should be as feasible as vaccination against α-HPV infections, which now are used to prevent anogenital and oropharyngeal cancers.

One possibility for the difficulty in detection of HPV in SCC is the low sensitivity and specificity of current whole tissue sequencing methods. To date, the application of single-cell technology for viral detection and viral genomics has been limited to low throughput and plate-based methods, including single cell genome and/or RNA sequencing to detect HIV DNA/RNA in isolated peripheral blood mononucleated single cells, (Josefsson et al., 2013; Wiegand et al., 2017), HPV18 in 40 HeLa S3 cells using a plate-based, modified SMART-seq2 protocol (Wu et al., 2015), and Hepatitis C virus using plate-based single-cell qPCR (McWilliam Leitch and McLauchlan, 2013). We propose that the recent advances in single-cell RNA-sequencing (scRNA-seq) technology can overcome these problems and enhance our ability to detect and associate the presence of HPV, and/or other viruses, with the development of cutaneous SCC. Importantly, scRNA-seq methods would also enable the identification of the specific cells expressing HPV transcripts, advancing our ability to understand the pathophysiology of HPV in SCC.

The K14E7 mouse model has been used to study the effect of the HPV *E7* viral transcript (Tuong et al., 2018). As proof-of-concept, we performed massively parallel scRNA-seq on keratinocytes isolated from the skin of K14E7 and control mice to show detection sensitivity and to provide insight into the effect of *E7* expression on transcription in basal cells. Our results provide evidence that HPV *E7* mRNA expression can be detected in K14E7 basal cells with high resolution (≥1 copy per cell) using high-throughput scRNA-seq. The approach presented here paves the way for the detection of HPV transcripts in human SCC of the skin, if present, and thereby provide insight into whether HPV is involved in the malignant progression of skin lesions.

## Results

### Whole tissue RNA-sequencing of human skin malignancies

To assess the association of HPV infection with human epithelial malignancy, we examined skin samples for the presence of HPV transcripts using standard RNA-sequencing technology. We performed standard paired-end RNA-sequencing on 25 skin samples diagnosed as healthy, AK, IEC or SCC collected from 17 individuals, and detected HPV transcripts in only one sample (Figure 1A), with reads mapping to multiple HPV genotypes, specifically HPV 96, 15 and 5 (Figure 1B). Most skin tumor samples were negative for HPV transcripts by this method. We hypothesized that if HPV infection was relevant to cancer initiation, HPV gene expression would likely be limited to basal keratinocytes, and possibly expressed at a low level, and thus, the heterogeneous composition of the tissue samples would reduce the effectiveness and sensitivity of detection by standard RNA-sequencing. These results led to examining the potential of scRNA-seq for detection of HPV transcripts in epithelial tissues.

**Figure 1:**
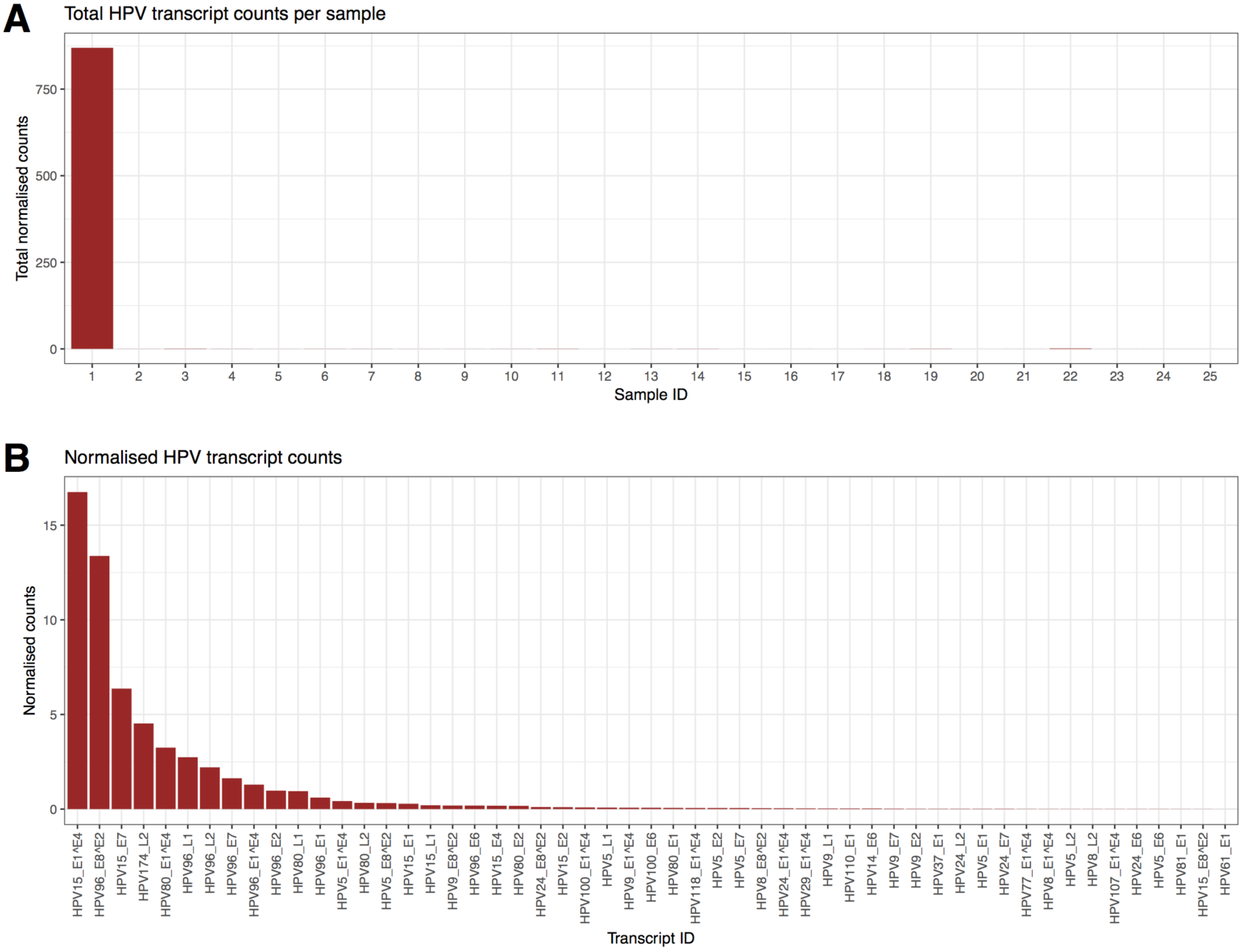
**Sensitivity of standard RNA-sequencing is insufficient to detect HPV transcription in the majority of human AK/SCC tissue samples.** (a) Total HPV transcript counts from standard whole-tissue RNA-sequencing were calculated for each of 25 samples from 17 individuals. Sample one, obtained from a shave biopsy from the forearm of an 81-year old female, is responsible for 99.3% of HPV expression observed in the cohort. (b) The median of the normalized HPV counts was obtained for each of the HPV transcripts with non-zero expression values and arranged by transcript abundance across the entire cohort. HPV splice variant IDs are marked with a ‘ˆ’ between the ORF IDs (e.g. HPV15_E1ˆE4).

### Single-cell RNA-sequencing of C57/BL6 and K14E7 mouse lesions

To determine whether scRNA-seq could detect expression of HPV genes in keratinocytes at the single-cell level, we examined skin from a mouse transgenic for the HPV16 *E7* oncogene driven from a Keratin 14 promoter (K14E7) (Lambert et al., 1993). We have previously described the use of this transgenic mouse to model pre-cancerous hyperproliferative epithelium (Tuong et al., 2018; Zhussupbekova et al., 2016). The scRNA-seq data were generated using the Chromium Single Cell 3’ RNA-sequencing method (10X Genomics) (Zheng et al., 2017a), which includes unique molecular identifier barcodes, enabling the copy number of detected transcript molecules to be determined. We obtained high quality sequence data for 9,410 single cells, of which 3,964 cells were from K14E7 epidermis and 5,446 from C57BL/6 epidermis. Following quality control (Methods) we achieved an average of 23,938 reads per cell, and detected expression of a median of 1,092 genes per cell.

### E7 mRNA is detectable in single K14E7 epidermal cells

We detected the expression of HPV16 *E7* transcripts in 832 (21%) of cells from the K14E7 cells, and, as expected, no C57BL/6 cells expressed HPV genes (Figure 2A). On average, *E7* positive cells expressed 1.93 *E7* mRNA transcripts/cell, and the maximum copy number observed was 14 transcripts in a single cell (Figure 2B). We subsequently investigated genome-wide differences in gene expression between the K14E7 and C57BL/6 cells using differential gene expression analysis. As anticipated, we observed a strong, statistically significant upregulation of *E7* in the K14E7 cells as a log2 fold-change of 5.65 (*p* = 2.94 × 10^−58^), which equates to a 50.4-fold linear increase in expression (Figure 2C). In total, we observed 140 significant differentially expressed genes (126 up-regulated; 14 down-regulated; adj. *p* < 0.01), which may be attributable to over-expression of the E7 transgene. Several of these gene changes (e.g. *E7*, *Krt16*, *Krt15*, *H2-Ab1*, *Cd74*, *Sprr1b* etc.) are concordant with our previous results from whole tissue RNA-seq (Tuong et al., 2018; Zhussupbekova et al., 2016). The results of this analysis are presented in Supplementary Table 1.

**Figure 2:**
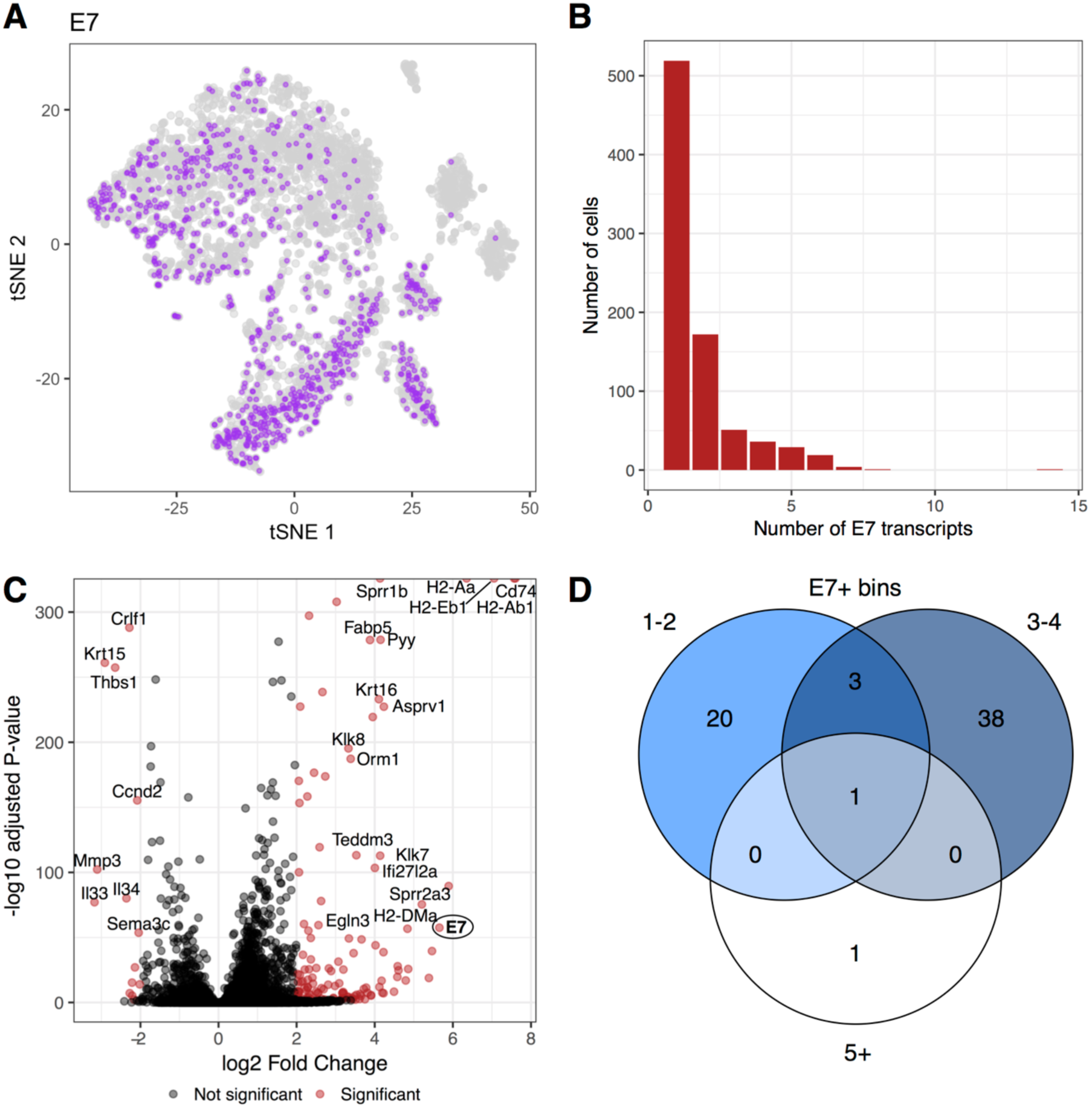
**HPV16-E7 transcript detection in single cells** (a) *t*-SNE visualization of *E7* positive and negative cells in transgenic samples. Each point represents a single cell reflecting the binary expression (on/off) of the *E7* transcript. Purple indicates the cell expresses ≥ 1 *E7* transcript, whereas the grey points represent zero *E7* expression in that cell. (b) Histogram showing the number of *E7* positive transcripts detected in transgenic cells. The x-axis shows the number of E7 UMI counts detected per cell and the y-axis indicates the number of cells with that number of E7 transcripts. (c) Differential expression between the wildtype and HPV16 E7 transgenic samples. The red points represent significant differentially expressed genes (adjusted *p* < 0.01) with a log2FC ≥ 2, whereas the black points represent the remaining non-significant genes. The E7 transcript is shown in bold and encircled. (d) Cells expressing *E7* were classified based on the number of *E7* transcripts detected and differential expression was performed between each bin and the remaining bins (e.g. bin A vs bin B+C). The Venn diagram shows the number of significant overlapping genes from each test.

### The effects of increasing E7 copy number on gene expression

Increased expression of *E7* mRNA per cell could impact upon the expression levels of other genes. We used two strategies to investigate at the single-cell level whether gene expression changes resulted from an increased expression of *E7* mRNA. First, we used linear regression to examine the effect of *E7* mRNA transcript number on the expression of each expressed gene (*m* = 13,744) in *E7* mRNA+ cells (*n* = 832). We identified 69 genes with statistically significant (false discovery rate < 0.1) changes related to *E7* mRNA transcript number. For all 69 genes, their expression levels increased with *E7* copy number. Notably, the genes with the largest effect of E7 copy number were *Krt14* (*β* = 10.54, *p* = 4.9×10^−4^) and *Krt5* (*β* = 1.79, *p* = 9.1×10^−5^), suggesting that basal keratinocytes are more likely to be influenced by increased *E7* mRNA expression. Summary statistics for all gene are presented in Supplementary Table 2.

Linear regression revealed a link between the number of *E7* transcripts per cell and the expression levels of multiple genes. We anticipated that there would be observable differences between the expression profiles of cells with low, medium and high numbers of *E7* transcripts. We used a differential expression approach to understand the differences between cells grouped by *E7* mRNA copy number. Here, we created bins of *E7*+ cells based on their *E7* mRNA copy number, such that bins contained cells with 1-2 (bin A), 3-4 (bin B), or 5+ copies (bin C). We examined differential gene expression each bin and the combined remaining bins to separate the transcriptome signature of each bin; bin A was compared to bins B+C, bin B to bins A+C and bin C to bins A+B. Our results showed that *E7* was the top differentially expressed gene in each analysis. We also noted that *Krt5* and *Krt14* were the next most significantly expressed in bin A cells compared to the remaining cells (log2 FC (*Krt5*) = 1.31, *p* = 1×10^−24^; log2 FC (*Krt14*) = 1.14, *p* = 5.7×10^−22^). In addition, our finding that increased E7 copy number is linked to increased keratin gene expression was further supported by our finding that cells in bin A (1-2 copies of E7) had a significantly lower expression mean for both genes than cells in bin B (3-4 copies of *E7*; mean *Krt5_binA_* = 15.56, mean *Krt5_binB_* = 20.46; t-test, p = 0.0451 and mean *Krt14_binA_* = 104.79, mean *Krt14_binB_* = 138.84; t-test, p = 0.0376). Genes that are differentially expressed across multiple *E7* bins may represent components of important biological processes linked to E7 transgene expression, compared to bin-specific genes. From the results of the differential expression analysis across bins, we observed an overlap of only 1 gene between all three bins (*E7*) and only 3 genes between bin A and B (*E7*, *Krt5*, *Krt14*) (Figure 2D), which is likely due to the differences in cell number per bin. The complete tables of significant differential expression results between bin groups are presented in Supplementary Tables 3-5. Thus, increased *E7* mRNA expression in single cells is predominantly associated with increased cellular expression of the basal keratins Krt5 and Krt14.

### Keratin/E7 co-expression in basal and supra-basal keratinocytes

We anticipated that the majority of cells expressing the K14E7 transgene would be basal keratinocytes, and as basal cells differentiate, the expression of *E7* mRNA would be reduced or lost. Cells expressing *Krt1* and *Krt10* represent more differentiated, supra-basal keratinocytes (Fuchs and Green, 1980). Hence, we determined which cells co-expressed *E7* and keratin mRNA transcripts. Figure 3A and B show a *t*-distributed stochastic neighbor embedding (*t*-SNE) figure to visualize cells that co-express important keratin subunits (*Krt5*/*Krt14* and *Krt1*/*Krt10*), while Figure 3C and D shows the cells that co-express the keratin subunits with *E7*. This analysis shows that while there is some specificity for basal keratinocytes, differentiating keratinocytes also expressed *E7* mRNA.

**Figure 3:**
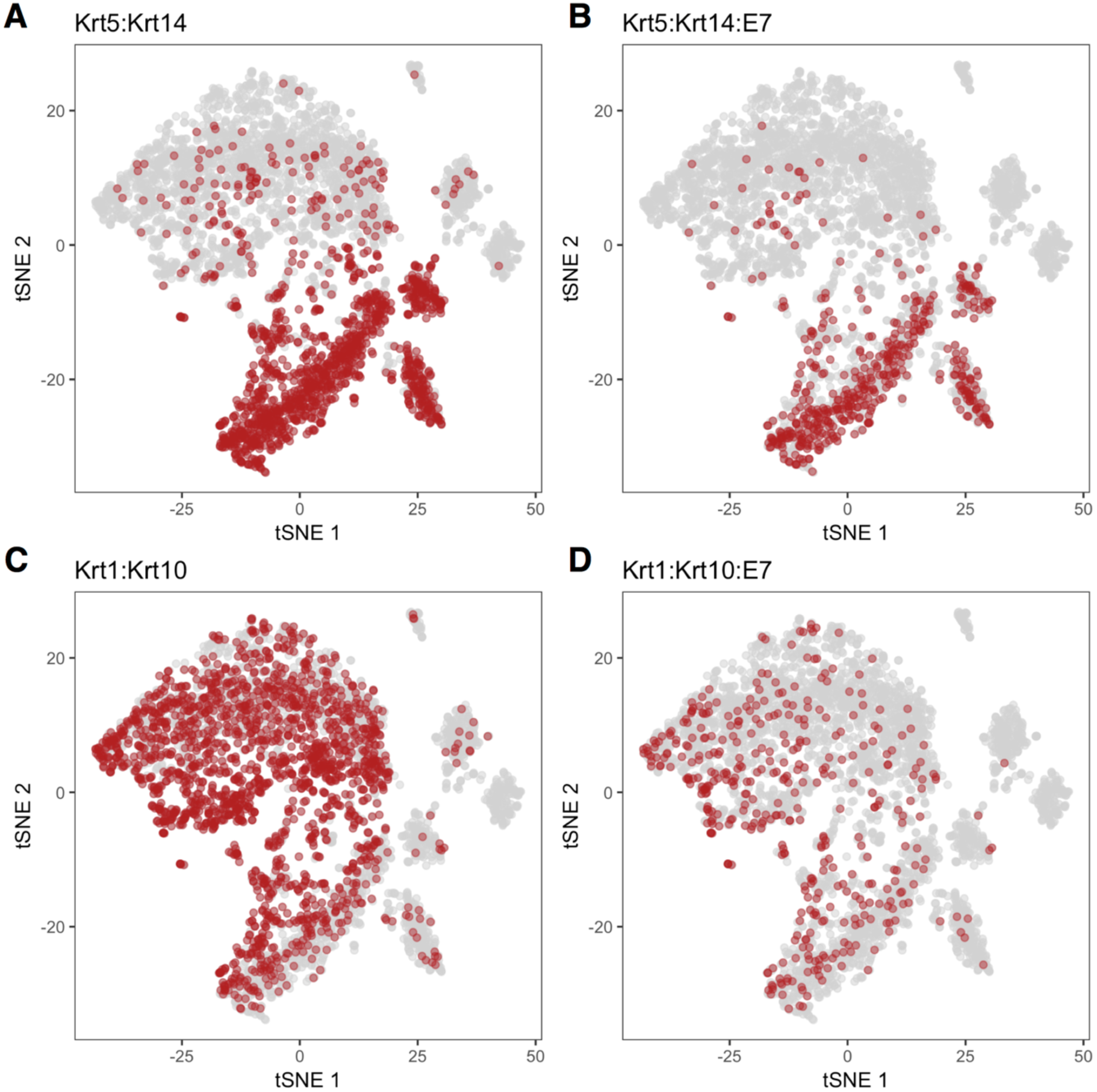
***E7* co-expression with major keratin genes** Each panel contains a *t*-SNE visualization of transgenic cells that co-express specific keratin genes (double-positive) that define basal or supra-basal cells (panels a and c), and those that also express E7 (triple-positive, panels b and d). (a) *Krt5* and *Krt14*, (b) *Krt5*, *Krt14* and *E7*, (c) *Krt1* and *Krt10*, and (d) *Krt1*, *Krt10* and *E7*.

To more clearly define which dermal layers harbor *E7* mRNA+ cells, we interrogated our data as a cohort of 832 individual *E7*+ transcriptomes, and correlated the expression of *E7* mRNA with each of the major keratin genes that define proliferating (*Krt14, Krt5*) or differentiated (*Krt1, Krt10*) keratinocytes. Using Spearman rank correlations, we calculated the correlation of *E7* mRNA expression between each of the major keratin subunits (*Krt14*, *Krt5*, *Krt10*, *Krt1*). Our results show that *E7* mRNA expression has a positive correlation with *Krt14* (Spearman’s *ρ* = 0.106, *p* = 2.3×10^−3^) and *Krt5* (Spearman’s *ρ* = 0.136, *p* = 8.15×10^−5^), and was negatively correlated with *Krt10* (Spearman’s *ρ* = -0.266, *p* = 5.91×10^−15^) and *Krt1* (Spearman’s *ρ* = -0.172, *p* = 6.08×10^−7^).

Overall, the data suggest that although *E7* is expressed in both proliferating and differentiating cell types, its effects on gene expression can be observed predominantly in basal keratinocytes. Conversely, *E7* co-expression with *Krt1*/*Krt10* in supra-basal keratinocytes is in the opposite direction, indicating *E7* expression is less effective in supra-basal cells and may be residual as keratinocytes become more differentiated.

### E7 is the major HPV16 gene expressed in K14E7 skin cells

Finally, to further confirm that the HPV16 *E7* transgene was the HPV-based driver of changes in the E7 transgenic cells, we examined the location of the mapped sequencing reads associated with the HPV16 genome. Since the *Krt14/E7* transgene also contains the *E6* gene, albeit with a transcription termination linker in the E6 DNA, we anticipated some reads would map to the *E6* region. Our results confirmed that *E7* is the predominant HPV16 gene expressed in the K14E7 mouse tissue (Figure 4A and B) although we also observed a small number of reads mapped to the *E6* region.

**Figure 4:**
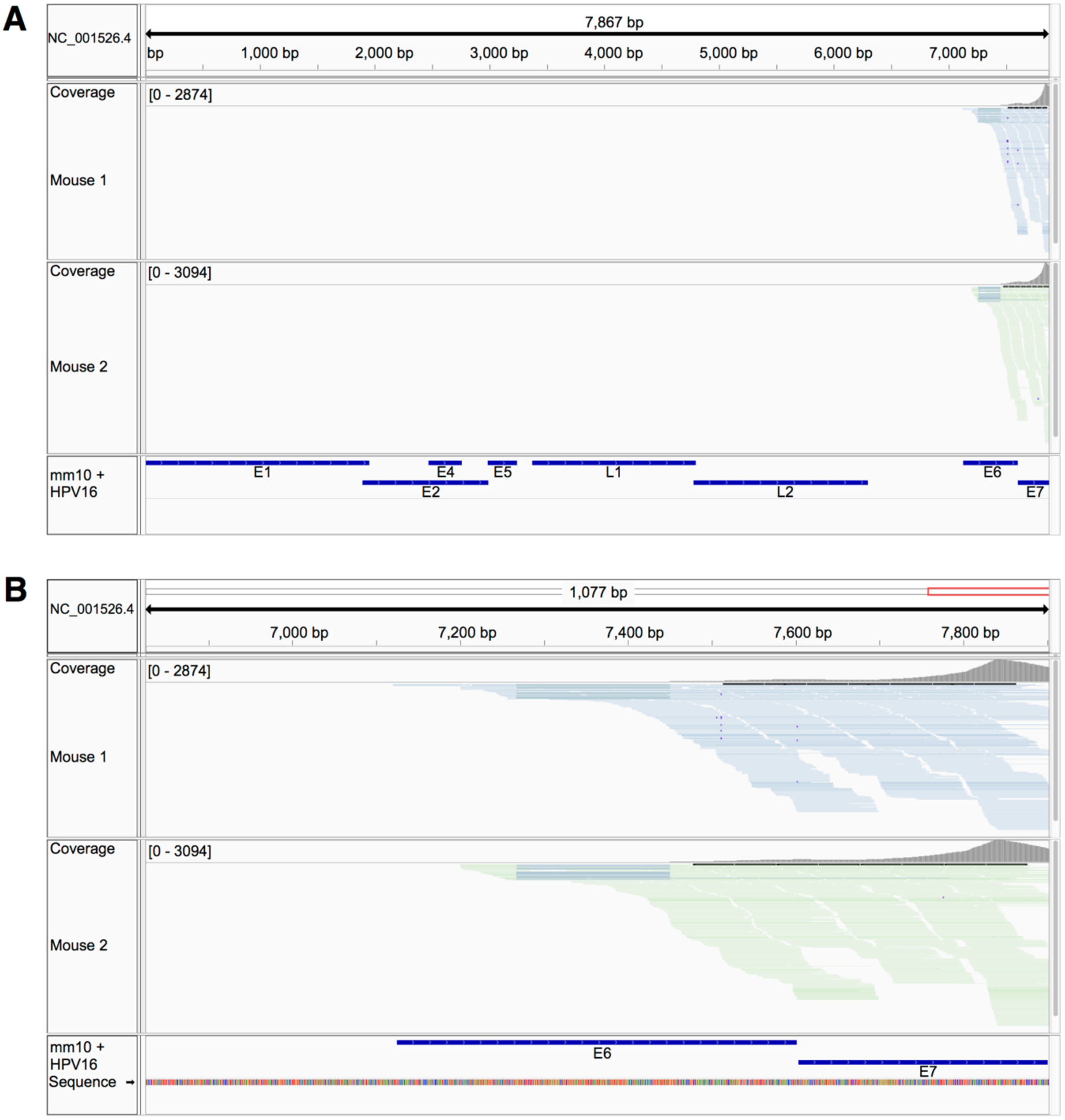
**Visualization of mapped E7 reads to the HPV16 genome** IGV visualization of sequencing reads from transgenic samples 1 (blue) and 2 (green) and their mapping specificity in the HPV16 genome. Panel a) shows the full length HPV16 genome and panel b) shows the immediate region surrounding the *E6* and *E7* loci. Evidence of splicing around the transcription termination linker present in the E6 gene is shown by the blue-grey bars in each between ∼7.25-7.45kb.

## Discussion

In this proof-of-concept study, we have demonstrated that massively parallel scRNA-seq can detect the presence of viral transcripts in a murine HPV16 transgenic mouse model. To our knowledge, this is the first time that HPV16 *E7* transcript expression has been captured and measured with high resolution, allowing us to examine the transcriptome-wide effects of HPV16 *E7* transgene expression in large numbers of individual cells.

The most commonly detected HPV DNA in non-melanoma skin cancer (NMSC) belong to those of the β-HPV subtypes. ‘High-risk’ β-HPVs (e.g. HPV5 and HPV8) are known to be particularly associated with EV lesions and have been implicated in cancer development in EV patients but not the general population (Gewirtzman et al., 2008). There is extensive literature reporting a variety of techniques that have been used to qualitatively, and in some instances (semi)-quantitatively, assess the presence of HPV in cutaneous tissues. In general, DNA genotyping appears to be the default method for assessing presence of HPV. These methods include (*i*) HPV viral DNA detection by PCR amplification of the conserved HPV L1 open reading frame or type-specific HPV E7 multiplex PCR and DNA microarray genotyping assays, (*ii*) PCR-amplification followed by reverse hybridization (e.g. Roche Linear Array HPV), (*iii*) a combination of DNA hybridization and chemiluminescent signal amplification (*e.g.* QIAGEN Digene Hybrid Capture 2 High-Risk DNA Test), PCR-amplification followed by Sanger sequencing, *in situ* hybridization of viral DNA. However, these methods have not conclusively ascertained whether a productive HPV replication cycle is required for, or involved in, cutaneous carcinogenesis, but showed only that a higher copy number of total HPV, rather than any specific type, is associated with cutaneous SCCs (Neale et al., 2013).

Indirect methods such as p16^INK4a^ staining (protein accumulation of CDKN2A) and detection of seroreactivity to HPV pseudovirions or HPV fusion proteins in peripheral blood can inform on HPV activity or productive presence, and have been used as a surrogate for classifying HPV positivity in patients. However, while there is a clear association of p16 accumulation with actinic keratosis progression to NMSC (Hodges and Smoller, 2002), they do not necessarily correlate with HPV DNA genotyping results (Svajdler et al., 2016) and the accuracy of the serology testing is debatable (Iannacone et al., 2014). Conversely, there are limited reports on the detection of HPV mRNA in cutaneous lesions, which we surmised to be most likely due to the inability, or lack of sensitivity of current methods, to detect the presence of the transcripts accurately. Interestingly, HPV16 E6 DNA and E6*I mRNA have been detected in 3/6 cases of primary squamous cell scrotal carcinoma using the QuantiTect Virus Kit and specific primers (Guimera et al., 2017). This might be due to the proximity to the anogenital region where HPV16 is most commonly located. Finally, mRNA sequencing of 31 cutaneous squamous cancers was unable to detect HPV transcripts even in samples that were HPV DNA positive (Arron et al., 2011). We reproduced this finding in a cohort of 25 AK/SCC/IEC patient lesions, and of these, we detected HPV transcripts in only 1 sample. This single sample, a shave biopsy from the forearm of an 81-year old female, accounted for 99.3% of HPV transcript counts, whereas the remaining samples were below the limits of detection. In particular, we detected transcripts corresponding to all capsid genes and early ORFs belonging to HPV96, and early ORFs for HPV15 in this sample. However, the presence of the transcripts encoding the capsid proteins belonging to HPV96 indicate that this was likely an active infection where the virus was undergoing a productive life cycle. The expression of the HPV5 early ORF in the same sample suggests that perhaps HPV15 transformed cells are permissive for vegetative reproduction of HPV96. Nevertheless, this is an isolated incident and is only speculative. Thus, whether HPV is associated with the transformation process in cutaneous NMSC remains unclear.

We have used the HPV 16 E7 mouse model to characterize the nature of the immune environment in hyperproliferative epithelium in detail by grafting *E7* transgenic and hyperproliferative epithelium to immune competent mice, and manipulating components of the immune system in either the graft recipient or the graft donor. We previously used this approach to establish that bone marrow-derived NKT cells recruited to hyperproliferative epithelium in the donor negatively regulate immune effector function by an IFN-γ dependent mechanism (Mattarollo et al., 2010). The hyperproliferative epithelium of the K14E7 mouse model also shares transcriptional characteristics of human AK premalignant lesions (Nindl et al., 2006). We note the caveat in the use of the transgenic HPV 16 (α-HPV) E7 mouse model for this analysis as it does not reflect the most likely subtype of HPV that may be found in human skin (β-HPVs). However, we anticipate that single cell sequencing technology, such as the method used here, should be amenable to detect all viral mRNA if the genes are transcribed.

One potential complication is the polycistronic nature of HPV mRNAs; RNAs encoding HPV genes are spliced to contain segments spanning multiple genes, resulting in different transcript variants that can encode different genes (McBride and Warburton, 2017). The spliced viral transcripts are generated with either the ‘early’ or ‘late’ polyadenylation sites following E5 and L1 respectively, a limitation that will be encountered by the 10X Chromium 3’ system, but one that would be overcome with full length scRNA-seq. In the K14E7 model, the transgene is engineered with the human Keratin 14 polyadenylation sequences immediately following the *E7* sequence (Herber et al., 1996), thereby allowing for transgenic expression of the *E7* gene, and thus enabling detection with the 10X Chromium 3’ system. Another potential challenging aspect is the processing steps required to generate enough viable cells for the sequencing; whilst preparation of viable single cell suspensions from murine epidermal sheets is a challenging process in and of itself, we anticipate that the difficulty will be even more pronounced when processing viable epidermal single cell suspensions from human skin biopsies. Much of this may be overcome with the use of single cell digital droplet PCR technology where only a small number of cells and input material is required for the assay. Indeed, digital droplet PCR technology has already been applied in diagnostic labs/clinics and for oropharyngeal squamous cancer where it was shown to be accurate for detecting HPV16 in the cancers (Biron et al., 2016). However, the 10X Chromium scRNA-seq platform remains an attractive option as it allows for high-throughput assessment of the mRNA transcriptome in tens of thousands of single cells, which we argue is needed to increase the overall sensitivity, both by cell number and the number of genes detected per cell, for detecting HPV(s) in single cells during skin carcinogenesis.

In this study, we have shown that (*i*) the expression of viral transcripts can be accurately measured with unprecedented resolution in single cells using high-throughput droplet-based technology, (*ii*) the transcriptome-wide effects of the viral transcripts can be observed and quantified in individual cells, (*iii*) increased *E7* transcript copy number is associated with increased expression of biologically relevant host genes, and (*iv*) E7 expression is predominantly associated with basal keratinocytes. In conclusion, the high-resolution quantitation afforded by droplet-based scRNA-seq will significantly enhance our understanding of the viral mechanisms of cancer progression, particularly those where the role of HPV is ambiguous, such as in oral cavity squamous cancers, and shows great promise for clinical diagnostic and prognostic applications.

## Acknowledgments

We thank the core facilities at the Translational Research Institute for excellent technical assistance in animal care (Biological Research Facility) and flow cytometric sorting (Flow Cytometry Core Facility, The sequencing Facility at the Institute for Molecular Bioscience, The University of Queensland, for sequencing the single cell RNA libraries, The University of Queensland Centre for Clinical Genomics at The University of Queensland Diamantina Institute for sequencing the whole tissue RNA-seq libraries. This work is funded from public competitive grant awarding bodies with no involvement in the conduct of the research, or production of the manuscript. Specifically, this work was supported by research project grants from the National Health and Medical Research Council (NHMRC), Merchant Charitable Foundation and Australasian College of Dermatologists Scientific Research Fund. Z.K.T. is supported by an Advance Queensland Early Career Research Fellowship, J.E.P. is supported by a NHMRC Career Development Fellowship and H.P.S. is supported by a NHMRC Practitioner Fellowship. No payment was received from any pharmaceutical company or other agency to write this manuscript.

## Author Contributions

S.W.L., Z.K.T., and K.N. conducted the experiments; S.W.L., Z.K.T. and A.S. analyzed the data; Q.H.N. and H.P.S. provided intellectual input; S.W.L., Z.K.T., K.N, I.H.F and J.E.P designed the experiments and wrote the paper.

## Materials and Methods

### Collection of human samples

Skin lesions and healthy skin tissue samples were collected from consenting patients that presented to the Dermatology department in Princess Alexandra Hospital from 2011-2013. The study was approved by Metro South Human Research Ethics Committee and The University of Queensland Human Research Ethics Committee (HREC-11-QPAH-236, HREC-11-QPAH-477, HREC-12-QPAH-217, and HREC-12-QPAH-25).

### E7 transgenic mouse model

C57BL/6 and K14E7 mice were maintained at the Translational Research Institute Biological Resources Facility (TRI BRF, Brisbane). All mice were maintained under pathogen free conditions and sex matched (all female). All procedures were approved by The University of Queensland Animal Ethics Committee (UQDI/367/13/NHMRC and UQDI/452/16).

### Tissue collection and dissociation

Ear skin from 10-12-week-old mice (*n*=2 per strain) was split into dorsal and ventral parts and incubated with 2.5 μg/μL Dispase II (Roche) for 1 h at 37°C. Epidermis and dermis were separated with closed forceps. The epidermis was further homogenized and digested with 1 μg/μL of collagenase D (Roche) and 0.2 μg/μL of DNase, (Roche) for 1 h at 37°C. Digested samples were passed through a 0.7 μm filter (BD Falcon) to generate a single cell suspension for staining and sorting.

### Flow cytometry

Single cell suspensions of digested epidermis samples were incubated with Fc Block (5 μg, rat anti-mouse CD19/CD32, clone 93, eBioscience) diluted in PBS for 30 min on ice. Samples were subsequently incubated with APC-conjugated rat anti-mouse CD45.2 antibodies (0.5 μg, Clone 104, Biolegend) diluted in PBS + 2% serum + 2 mM EDTA for 30 min on ice. Prior to sorting, cells were labelled with 7-AAD (0.25 μg, eBioscience). Live CD45‐ cells (7-AAD‐ and APC‐ or PE-Cy7-) were sorted using the BD ARIA Fusion sorter at 12 psi with a 100 μm nozzle. ∼300,000-500,000 events/cells were collected per sample.

### Whole-tissue RNA-sequencing

Total RNA was extracted using TRIzol (Life Technologies) and RNA purification was performed using the RNeasy mini kit (Qiagen). RNA was assessed for quality and quantity using a Nanodrop instrument (ThermoFisher Scientific). RNA sequencing libraries of poly (A) RNA from 500ng total RNA obtained from AK, IEC and SCC skin lesions were generated using the TruSeq unstranded mRNA library prep kit for Illumina multiplexed sequencing (Illumina). Libraries were sequenced (100 base pair, paired-end) on the Illumina HiSeq 2500 platform (Illumina).

### Single cell RNA-sequencing (scRNA-seq)

scRNA-seq was performed in duplicate for wild-type and K14E7 transgenic animals. The 10X Genomics Chromium instrument (10X Genomics) was used to partition viable CD45- cells with barcoded beads, and cDNA from each cell was prepared using the Single Cell 3’ Library, Gel Bead and Multiplex Kit (v1; 10X Genomics; PN-120233) as per the manufacturer’s instructions. Cell numbers in each reaction were optimized to capture approximately 3,000 cells. cDNA shearing was performed with a Covaris S2 instrument (Covaris) set to produce a target size of 200bp (Intensity:5, Duty cycle: 10%; Cycles: 200; Time: 120s). The resulting single cell transcriptome libraries were pooled and sequenced on an Illumina NextSeq500, using a 150-cycle High Output reagent kit (NextSeq500/550 v2; Illumina, FC-404-2002) in standalone mode as follows: 98bp (Read 1), 14bp (I7 Index), 8bp (I5 Index), and 10bp (Read 2).

### Bioinformatics processing

Whole-tissue RNA-sequence reads were mapped to a custom reference containing the human GRCh38 transcriptome and alpha and beta HPV CDS and spliced sequences with Salmon v.0.9.1 (Patro et al., 2017) using the default parameters. HPV sequences were downloaded from pave.niaid.nih.gov (Van Doorslaer et al., 2017). Length-scaled transcript per million (TPM) values for each transcript were normalized for library size differences using edgeR v3.16.5 (Robinson et al., 2010), and, to attain maximum sensitivity, transcript counts were retained if there was at least 1 count in 1 sample.

For the single-cell data, the *cellranger* pipeline v1.3.1 provided by 10X Genomics (*mkfastq*, *count*, *aggr*) (Zheng et al., 2017b) was used to process the raw sequencing data using the default parameters, and adjusted to the expected number of cells per sample (3,000). Reads were aligned to a custom reference genome comprising the mouse (mm10) and the HPV16 genome (NC_001526) using the STAR aligner (Dobin et al., 2013) included in the *cellranger* pipeline. Quality control of cell barcodes and unique molecular identifiers was performed during the *cellranger count* stage using default parameters. A between-sample normalized gene expression matrix for four samples was generated using *cellranger aggr* for further analysis.

### Single cell RNA-sequence analysis

The aggregated single cell gene expression data generated by *cellranger* was used as the input for the *ascend* workflow (Senabouth et al., 2017). Expression levels for each transcript were determined using the number of unique molecular identifiers (UMI) assigned to the transcript. The data was imported into an EMSet object using the LoadCellRanger function. Quality control and filtering steps were performed to remove outlier cells and genes. Cells were removed if the library size, or number of expressed genes exceeded 3 median absolute deviations, and also if a cell expressed high proportions of mitochondrial (>20%) or ribosomal (>50%) genes. Following cell-cell normalization by relative log expression (RLE), ribosomal and mitochondrial control genes were removed from the analysis.

Principal component analysis was performed on the filtered and normalized gene expression matrix, and the first 20 PCs that explained the majority of variance in the data were retained. To visualize the patterns of gene expression in each cell, *t*-distributed stochastic neighbor (*t*-SNE) projections were generated using the PCA-reduced data.

Differential expression analysis was performed with DESeq (Anders and Huber, 2010) in *ascend* between (i) wildtype and HPV16 E7 transgenic cells and (ii) bins of E7 positive cells, where bins of cells containing 1-2, 3-4 or 5+ copies of *E7* were generated. Genes were considered significant if the adjusted p-value was below the multiple-testing threshold of 0.01 (Benjamini-Hochberg method) and the absolute log2 expression fold change was ≥ 2.

Position-sorted BAM files for the transgenic samples were generated by the *cellranger* software pipeline and these were used for visualization of mapped reads using the IGV software (Robinson et al., 2011).

### Linear modelling of transcript expression

The expression profile of retained transcripts was tested for significant association with HPV16 *E7* expression using linear regression. *E7*-expressing cells were extracted (*n* = 832) and 13,744 genes with non-zero expression were retained. Linear regression modelling tested the expression of each individual gene against the bins of cells with increasing *E7* copy numbers. The overall model *p*-value was corrected for multiple testing using the Benjamini-Hochberg method (FDR) and transcripts with an adjusted p-value below the FDR threshold (<0.1) were considered statistically significant.

### Data availability

RNA sequencing datasets were deposited in ArrayExpress. Single cell RNA-seq data is available via: https://www.ebi.ac.uk/arrayexpress/experiments/E-MTAB-6429/. Whole tissue RNA-seq data is available via: https://www.ebi.ac.uk/arrayexpress/experiments/E-MTAB-6430/.

## References

Anders, S., and Huber, W. (2010). Differential expression analysis for sequence count data. Genome Biol 11, R106.

Arron, S.T., Ruby, J.G., Dybbro, E., Ganem, D., and Derisi, J.L. (2011). Transcriptome sequencing demonstrates that human papillomavirus is not active in cutaneous squamous cell carcinoma. J Invest Dermatol 131, 1745–1753.

Biron, V.L., Kostiuk, M., Isaac, A., Puttagunta, L., O’Connell, D.A., Harris, J., Cote, D.W., and Seikaly, H. (2016). Detection of human papillomavirus type 16 in oropharyngeal squamous cell carcinoma using droplet digital polymerase chain reaction. Cancer 122, 1544–1551.

Boukamp, P. (2005). Non-melanoma skin cancer: what drives tumor development and progression? Carcinogenesis 26, 1657–1667.

Criscione, V.D., Weinstock, M.A., Naylor, M.F., Luque, C., Eide, M.J., Bingham, S.F., and Department of Veteran Affairs Topical Tretinoin Chemoprevention Trial, G. (2009). Actinic keratoses: Natural history and risk of malignant transformation in the Veterans Affairs Topical Tretinoin Chemoprevention Trial. Cancer 115, 2523–2530.

Dobin, A., Davis, C.A., Schlesinger, F., Drenkow, J., Zaleski, C., Jha, S., Batut, P., Chaisson, M., and Gingeras, T.R. (2013). STAR: ultrafast universal RNA-seq aligner. Bioinformatics 29, 15–21.

Farzan, S.F., Waterboer, T., Gui, J., Nelson, H.H., Li, Z., Michael, K.M., Perry, A.E., Spencer, S.K., Demidenko, E., Green, A.C., et al. (2013). Cutaneous alpha, beta and gamma human papillomaviruses in relation to squamous cell carcinoma of the skin: a population-based study. Int J Cancer 133, 1713–1720.

Feldman, S.R., and Fleischer, A.B., Jr. (2011). Progression of actinic keratosis to squamous cell carcinoma revisited: clinical and treatment implications. Cutis 87, 201–207.

Foote, J.A., Harris, R.B., Giuliano, A.R., Roe, D.J., Moon, T.E., Cartmel, B., and Alberts, D.S. (2001). Predictors for cutaneous basal‐ and squamous-cell carcinoma among actinically damaged adults. Int J Cancer 95, 7–11.

Forslund, O., Lindelof, B., Hradil, E., Nordin, P., Stenquist, B., Kirnbauer, R., Slupetzky, K., and Dillner, J. (2004). High prevalence of cutaneous human papillomavirus DNA on the top of skin tumors but not in “Stripped” biopsies from the same tumors. J Invest Dermatol 123, 388–394.

Franceschi, S., Levi, F., Randimbison, L., and La Vecchia, C. (1996). Site distribution of different types of skin cancer: new aetiological clues. Int J Cancer 67, 24–28.

Fuchs, E., and Green, H. (1980). Changes in keratin gene expression during terminal differentiation of the keratinocyte. Cell 19, 1033–1042.

Genders, R.E., Mazlom, H., Michel, A., Plasmeijer, E.I., Quint, K.D., Pawlita, M., van der Meijden, E., Waterboer, T., de Fijter, H., Claas, F.H., et al. (2015). The presence of betapapillomavirus antibodies around transplantation predicts the development of keratinocyte carcinoma in organ transplant recipients: a cohort study. J Invest Dermatol 135, 1275–1282.

Gewirtzman, A., Bartlett, B., and Tyring, S. (2008). Epidermodysplasia verruciformis and human papilloma virus. Curr Opin Infect Dis 21, 141–146.

Gordon, L.G., Elliott, T.M., Olsen, C.M., Pandeya, N., and Whiteman, D.C. (2017). Multiplicity of skin cancers in Queensland and their cost burden to government and patients. Aust N Z J Public Health.

Gordon, L.G., Scuffham, P.A., van der Pols, J.C., McBride, P., Williams, G.M., and Green, A.C. (2009). Regular sunscreen use is a cost-effective approach to skin cancer prevention in subtropical settings. J Invest Dermatol 129, 2766–2771.

Guimera, N., Alemany, L., Halec, G., Pawlita, M., Wain, G.V., Vailen, J.S.S., Azike, J.E., Jenkins, D., de Sanjose, S., Quint, W., et al. (2017). Human papillomavirus 16 is an aetiological factor of scrotal cancer. Br J Cancer 116, 1218–1222.

Herber, R., Liem, A., Pitot, H., and Lambert, P.F. (1996). Squamous epithelial hyperplasia and carcinoma in mice transgenic for the human papillomavirus type 16 E7 oncogene. J Virol 70, 1873–1881.

Hodges, A., and Smoller, B.R. (2002). Immunohistochemical comparison of p16 expression in actinic keratoses and squamous cell carcinomas of the skin. Mod Pathol 15, 1121–1125.

Iannacone, M.R., Gheit, T., Pfister, H., Giuliano, A.R., Messina, J.L., Fenske, N.A., Cherpelis, B.S., Sondak, V.K., Roetzheim, R.G., Silling, S., et al. (2014). Case-control study of genus-beta human papillomaviruses in plucked eyebrow hairs and cutaneous squamous cell carcinoma. Int J Cancer 134, 2231–2244.

Josefsson, L., Palmer, S., Faria, N.R., Lemey, P., Casazza, J., Ambrozak, D., Kearney, M., Shao, W., Kottilil, S., Sneller, M., et al. (2013). Single cell analysis of lymph node tissue from HIV-1 infected patients reveals that the majority of CD4+ T-cells contain one HIV-1 DNA molecule. PLoS Pathog 9, e1003432

Lambert, P.F., Pan, H., Pitot, H.C., Liem, A., Jackson, M., and Griep, A.E. (1993). Epidermal cancer associated with expression of human papillomavirus type 16 E6 and E7 oncogenes in the skin of transgenic mice. Proc Natl Acad Sci U S A 90, 5583–5587.

Madeleine, M.M., Patel, N.S., Plasmeijer, E.I., Engels, E.A., Bouwes Bavinck, J.N., Toland, A.E., Green, A.C., and the Keratinocyte Carcinoma Consortium Immunosuppression Working, G. (2017). Epidemiology of keratinocyte carcinomas after organ transplantation. Br J Dermatol 177, 1208–1216.

Mattarollo, S.R., Rahimpour, A., Choyce, A., Godfrey, D.I., Leggatt, G.R., and Frazer, I.H. (2010). Invariant NKT cells in hyperplastic skin induce a local immune suppressive environment by IFN-gamma production. J Immunol 184, 1242–1250.

McBride, A.A., and Warburton, A. (2017). The role of integration in oncogenic progression of HPV-associated cancers. PLoS Pathog 13, e1006211

McWilliam Leitch, E.C., and McLauchlan, J. (2013). Determining the cellular diversity of hepatitis C virus quasispecies by single-cell viral sequencing. J Virol 87, 12648–12655.

Neale, R.E., Weissenborn, S., Abeni, D., Bavinck, J.N., Euvrard, S., Feltkamp, M.C., Green, A.C., Harwood, C., de Koning, M., Naldi, L., et al. (2013). Human papillomavirus load in eyebrow hair follicles and risk of cutaneous squamous cell carcinoma. Cancer Epidemiol Biomarkers Prev 22, 719–727.

Patro, R., Duggal, G., Love, M.I., Irizarry, R.A., and Kingsford, C. (2017). Salmon provides fast and bias-aware quantification of transcript expression. Nat Methods 14, 417–419.

Pfister, H. (2003). Chapter 8: Human papillomavirus and skin cancer. J Natl Cancer Inst Monogr, 52–56.

Robinson, J.T., Thorvaldsdottir, H., Winckler, W., Guttman, M., Lander, E.S., Getz, G., and Mesirov, J.P. (2011). Integrative genomics viewer. Nat Biotechnol 29, 24–26.

Robinson, M.D., McCarthy, D.J., and Smyth, G.K. (2010). edgeR: a Bioconductor package for differential expression analysis of digital gene expression data. Bioinformatics 26, 139–140.

Senabouth, A., Lukowski, S.W., Alquicira, J., Andersen, S.B., Mei, X., Nguyen, Q.H., and Powell, J.E. (2017). ascend: R package for analysis of single cell RNA-seq data. bioRxiv.

Svajdler, M., Jr., Mezencev, R., Kaspirkova, J., Kacerovska, D., Kazakov, D.V., Ondic, O., and Michal, M. (2016). Human papillomavirus infection and p16 expression in the immunocompetent patients with extragenital/extraungual Bowen’s disease. Diagn Pathol 11, 53

Torre, L.A., Bray, F., Siegel, R.L., Ferlay, J., Lortet-Tieulent, J., and Jemal, A. (2015). Global cancer statistics, 2012. CA Cancer J Clin 65, 87–108.

Tuong, Z.K., Noske, K., Kuo, P., Bashaw, A.A., Teoh, S., and Frazer, I.H. (2018). Murine HPV16 E7-expressing transgenic skin effectively emulates the cellular and molecular features of human high-grade squamous intraepithelial lesions. Papillomavirus Research 5, 6–20.

Van Doorslaer, K., Li, Z., Xirasagar, S., Maes, P., Kaminsky, D., Liou, D., Sun, Q., Kaur, R., Huyen, Y., and McBride, A.A. (2017). The Papillomavirus Episteme: a major update to the papillomavirus sequence database. Nucleic Acids Res 45, D499–D506

Wheller, L., and Soyer, H.P. (2015). Clinical features of actinic keratoses and early squamous cell carcinoma. Curr Probl Dermatol 46, 58–63.

Wiegand, A., Spindler, J., Hong, F.F., Shao, W., Cyktor, J.C., Cillo, A.R., Halvas, E.K., Coffin, J.M., Mellors, J.W., and Kearney, M.F. (2017). Single-cell analysis of HIV-1 transcriptional activity reveals expression of proviruses in expanded clones during ART. Proc Natl Acad Sci U S A 114, E3659–E3668

Wu, L., Zhang, X., Zhao, Z., Wang, L., Li, B., Li, G., Dean, M., Yu, Q., Wang, Y., Lin, X., et al. (2015). Full-length single-cell RNA-seq applied to a viral human cancer: applications to HPV expression and splicing analysis in HeLa S3 cells. Gigascience 4, 51

Yu, S.H., Bordeaux, J.S., and Baron, E.D. (2014). The immune system and skin cancer. Adv Exp Med Biol 810, 182–191.

Zalaudek, I., Giacomel, J., Schmid, K., Bondino, S., Rosendahl, C., Cavicchini, S., Tourlaki, A., Gasparini, S., Bourne, P., Keir, J., et al. (2012). Dermatoscopy of facial actinic keratosis, intraepidermal carcinoma, and invasive squamous cell carcinoma: a progression model. J Am Acad Dermatol 66, 589–597.

Zheng, G.X., Terry, J.M., Belgrader, P., Ryvkin, P., Bent, Z.W., Wilson, R., Ziraldo, S.B., Wheeler, T.D., McDermott, G.P., Zhu, J., et al. (2017a). Massively parallel digital transcriptional profiling of single cells. Nat Commun 8, 14049

Zheng, G.X.Y., Terry, J.M., Belgrader, P., Ryvkin, P., Bent, Z.W., Wilson, R., Ziraldo, S.B., Wheeler, T.D., McDermott, G.P., Zhu, J., et al. (2017b). Massively parallel digital transcriptional profiling of single cells. Nat Commun 8, 14049

Zhussupbekova, S., Sinha, R., Kuo, P., Lambert, P.F., Frazer, I.H., and Tuong, Z.K. (2016). A Mouse Model of Hyperproliferative Human Epithelium Validated by Keratin Profiling Shows an Aberrant Cytoskeletal Response to Injury. EBioMedicine 9, 314–323.

